# MERISTEM-DEFECTIVE / DEFECTIVELY ORGANIZED TRIBUTARIES2 regulates the balance between stemness and differentiation in the root meristem through RNA splicing control

**DOI:** 10.1101/2022.11.23.517632

**Authors:** Helen L. Thompson, Weiran Shen, Rodrigo Matus, Medhavi Kakkar, Carl Jones, David Dolan, Sushma Grellscheid, Xiyan Yang, Na Zhang, Sina Mozaffari-Jovin, Chunli Chen, Xianlong Zhang, Jennifer F. Topping, Keith Lindsey

## Abstract

Plants respond to environmental stresses through controlled stem cell maintenance and meristem activity. One level of transcriptional control is RNA alternative splicing. However the mechanistic link between stress, meristem function and RNA splicing is poorly understood. The *MERISTEM-DEFECTIVE* (*MDF*)/*DEFECTIVELY ORGANIZED TRIBUTARIES* (*DOT2*) gene of Arabidopsis encodes a SR-related family protein, required for meristem function and leaf vascularization, and is the likely orthologue of the human SART1 and yeast snu66 splicing factors. MDF is required for the correct splicing and expression of key transcripts associated with root meristem function. We identified *RSZ33* and *ACC1*, both known to regulate cell patterning, as splicing targets required for MDF function in the meristem. *MDF* expression is modulated by osmotic and cold stress, associated with differential splicing and specific isoform accumulation and shuttling between nucleus and cytosol, and acts in part via a splicing target *SR34*. We propose a model in which MDF controls splicing in the root meristem to promote stemness and repress stress response and cell differentiation pathways.

**Summary statement:** The protein MERISTEM-DEFECTIVE regulates Arabidopsis meristem function through its role as a splicing factor, mediated through splicing targets RSZ33, ACC1 and SR34.

## INTRODUCTION

There is much interest in understanding the molecular mechanisms regulating meristem function, and interacting networks involving hormonal crosstalk and regulatory transcription factors have been identified (Santuari et al., 2016). Less well understood are the mechanisms determining how stress responses impact the balanced relationship between the stemness of meristems and cell differentiation processes, in order to control growth and development.

There is currently a large research effort into the role of RNA processing (microRNAs; long non-coding RNAs; nonsense-mediated mRNA decay, NMD; alternative splicing, AS) in the control of plant development and stress responses (Tanabe et al., 2006; Duque, 2011; Chen et al., 2013). There is good evidence for the role of AS in plant stress responses (Duque 2011), including temperature stress (Capovilla et al. 2018; Huertas et al. 2019), salt and nutrient stress (Lee et al. 2006; Nishida et al. 2017), response to abscisic acid (ABA; Zhu et al. 2017) or other environmental cues such as circadian rhythm (Cui et al. 2017); and in plant development (Szakonyi and Duque, 2018). However, while more than 300 genes encoding proteins putatively involved in splicing have been identified in Arabidopsis by homology searching (Reddy et al. 2013), very few have been tested experimentally for tissue-specific activities or in relation to the control of gene expression in response to external factors.

We previously identified the *MDF* gene (At5g16870) in an expression screen in the developing Arabidopsis embryonic root (Casson et al., 2005). It encodes a predicted arginine-serine (RS) domain protein, with homology to the human hSART-1 and yeast Snu66 proteins, which are splicing factors (Casson et al., 2009; Fig. S1). *MDF* is expressed at relatively high levels in the embryonic root meristem, and subsequently in the seedling root and shoot meristems, with lower levels of expression in vascular tissues (Fig. S1; Casson et al., 2005, 2009). It was also identified separately in a screen for vascular tissue-defective mutants, and given the name *DEFECTIVELY ORGANIZED TRIBUTARIES* (*DOT2*; Petricka et al., 2008). *mdf/dot2* loss-of-function mutants are dwarfed and both roots and shoot meristems show abnormalities, with defective root radial pattern (Fig. 1; Fig. S2). A very recent paper has shown that MDF is a likely splicing factor with a similar function to hSART-1 and mediates responses to DNA damage (de Luxan-Hernandez et al., 2022). Here we show that MDF is itself spliced, different isoforms are differentially expressed under cold and osmotic stress, and its function on cell patterning is mediated at least in part through control of splicing of the meristem regulators *RSZ33* and *ACC1*.

**Fig. 1.**
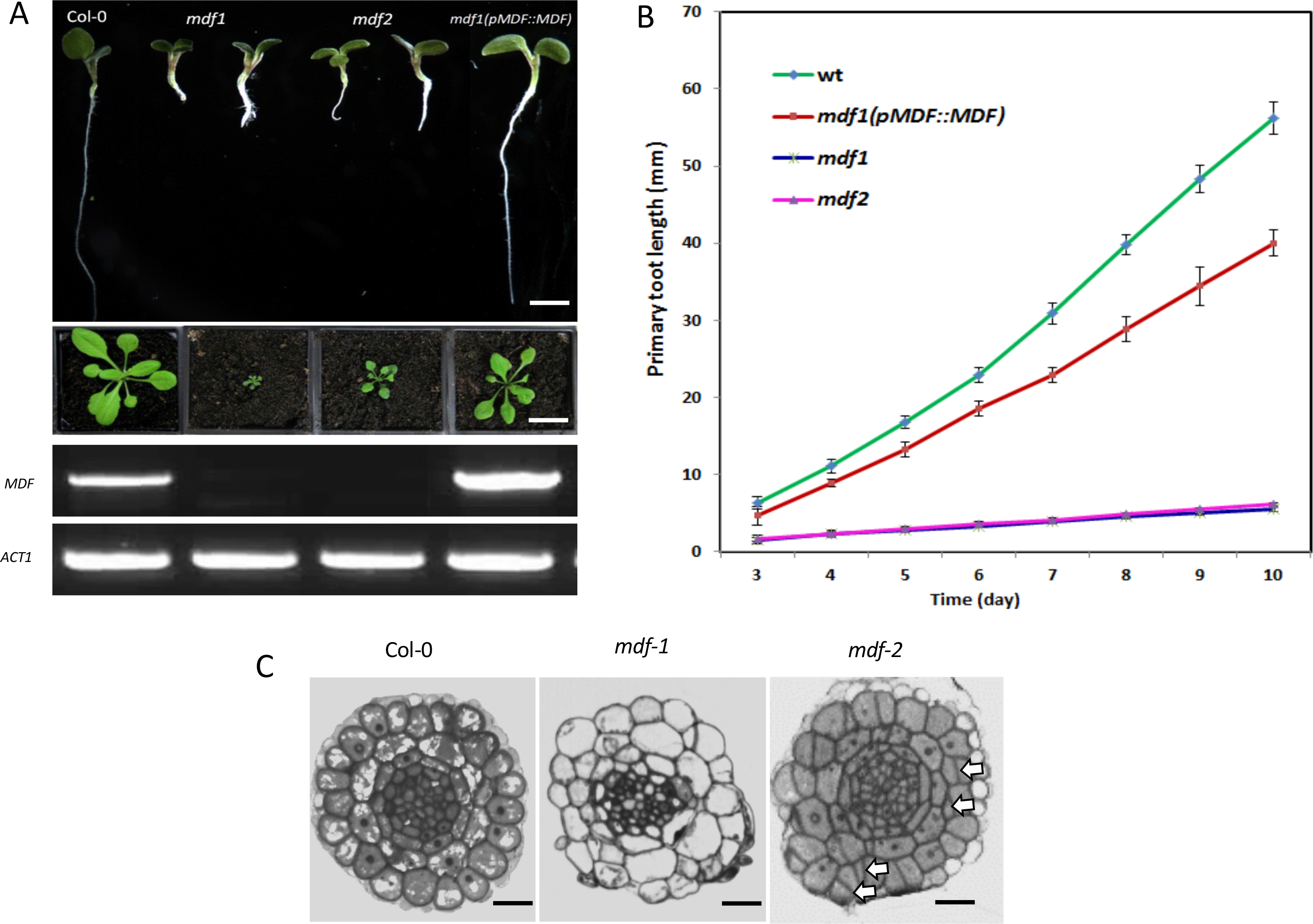
*mdf* mutants show defective cell division and growth. A. Upper panel: 10-day old seedlings of the two T-DNA mutants *mdf-1* and *mdf-2*, wild type Col-0 and a complemented *mdf-1* x *pMDF∷MDF* line. The more severe *mdf-1* mutant phenotype displayed stunted growth, short roots and was sterile, dying before 25 days. The *pMDF∷MDF* complementation of *mdf-1* largely rescued the mutant phenotype, following transformation of heterozygotes and selection of homozygote progeny. Scale bar = 0.5 cm. Middle panel: Phenotypes of wild type Col-0, *mdf-1*, *mdf-2* and the complemented *pMDF∷MDF* at 21 days post germination. Scale bar = 1.5 cm. Lower panel: Semi-quantitive RT-PCR of *MDF* in wild-type, both mutants and the complemented line. The full-length transcript was absent in both mutants. Control PCR with Actin primers ACT1 shown below. B. Change in primary root length over time for wild type Col-0, *mdf-1* and *mdf-2* mutants and the complemented line (*pMDF∷MDF*). C. Transverse sections through the root tip of wild type Col-0 *mdf-1* and *mdf-2* mutants. In Col-0 the boundaries between the cell layers are well defined. *mdf-1* has irregularly shaped cells in the epidermis, cortex and endodermis, and the boundary between the cell layers is poorly defined. *mdf-2* cells are more organized than in *mdf-1*, but show evidence of irregular divisions (arrows indicate some examples). Scale bar = 10 um.

## RESULTS AND DISCUSSION

### MDF/DOT2 is a component of the plant spliceosome

The predicted 820 amino acids MDF/DOT2 protein shares 41% identity with hSART-1 (Fig. S1). hSART-1 and the yeast homologue Snu66 are key components of the maturing spliceosome, involved in recruiting the tri-snRP during B complex formation, with secondary roles in cell cycle control (Makarova et al. 2001). Modelling of the putative 3-dimensional structure of the MDF/DOT2 protein suggests a strong structural similarity with hSART-1 (Fig. S2A). de Luxan-Hernandez et al. (2022) observed interaction between MDF and the spliceosome component PRP6 (also known as the plant homologue STABILIZED 1, STA1) though not with another tri-snRNP complex protein LSM8. To further investigate the possible molecular relationship between MDF/DOT2 and other splicing components in Arabidopsis, we carried out protein-protein interaction studies *in planta* with putative homologous plant spliceosome components Brr2a, PRP6 and PRP8A (Fig. S2B-I). While we found no detectable interaction between MDF/DOT2 and or PRP8A in BiFC (Fig. S2E) or in yeast-2 hybrid (not shown) experiments, we observed interaction with BRR2a, a splicing factor conserved in eukaryotes (Fig. S2 F, G; Mahrez et al., 2016); and with BIN2 (Fig. S2C), a nuclear-localized kinase required for root meristem maintenance that also interacts with the nuclear protein BZR1 required for root development (Fig. S2D; Yan et al., 2009; Li et al., 2020). These data strongly support the view that MDF/DOT2 is a structural component of the plant spliceosome complex.

### MDF promotes stemness and represses differentiation pathways

A role for MDF/DOT2 in splicing was further confirmed by RNA-sequencing on two independent *mdf* loss-of-function mutants (*mdf-1, mdf-2*; Casson et al., 2009; de Luxan-Hernandez et al., 2022) compared to wildtype. This analysis provided additional information on the biological pathways in which MDF plays a role, and also compared patterns of splicing isoform profiles in comparison with a wildtype control by analysing the data output from RMAts for alternative splicing (AS) events. We hypothesized that, if MDF/DOT2 acts as an essential splicing factor, we would see both altered patterns of RNA isoforms for some genes, and changes in the abundance of other, non-spliced, transcripts that may be targets of mis-spliced regulatory RNAs.

de Luxan-Hernandez et al. (2022) found upregulation of stress-related genes and down-regulation of cell division genes, among others. We found that 4195 genes were upregulated and 5404 genes downregulated in *mdf-1*, and 2830 were up and 3449 down in *mdf-2*, which has a less severe mutant phenotype after the seedling stage (Fig. 1A; Fig. S3A,B; Fig. S4A). Unbiased gene ontology (GO) enrichment analysis was carried out and treemaps were generated using data from REVIGO (Supek *et al*., 2011). Both *mdf-1* and *mdf-2* transcriptomes exhibit upregulation of stress-related genes (Fig. S4B-F), and enriched GO terms include ‘response to stress’, ‘defence response’, ‘response to other organism’, ‘immune response’, ‘reactive oxygen species metabolism’ and ‘response to ethylene’. In both tree maps, ‘aging’ and ‘programmed cell death’ were also shown to be significantly upregulated. This suggests that MDF plays a negative regulatory role in different stress response pathways.

The majority of the downregulated GO terms are related to signalling and development, including ‘auxin signalling pathway’, ‘protein phosphorylation’, and ‘tissue development and growth’. We previously showed altered auxin distribution in *mdf* mutant root tips, associated with defective PIN protein localization (which is dependent on correct *PLT* expression; Casson et al. 2009; Santuari et al., 2016; Aida et al. 2004) and potentially accounting for misexpression of auxin-regulated *WOX5* (Sarkar et al., 2007). The RNA-seq data show that the expression of several known root meristem regulation genes, including *PIN-FORMED* (*PIN1* to *PIN7*), *PLETHORA* (*PLT1*, *PLT2* and *PLT4*), *WUSCHEL-RELATED HOMEOBOX* (*WOX1*, *WOX4* and *WOX5*), *SHORTROOT* (*SHR*) and *POLARIS* are all significantly downregulated in *mdf-1*, and most also in the less severe *mdf-2*; *PLT5* is upregulated (Fig. S4G; Tables S1, S2). The radial pattern of the *mdf* primary root is abnormal (Fig. 1D), with similar supernumerary cell divisions to those seen in the *shr* mutant (Helariutta et al. 2000). Therefore MDF function is required for the correct expression and biological activity of a number of essential meristem genes. The PLT and SCR/SCR pathways that contribute to stem cell niche formation function independently (Aida et al. 2004), but our results show that both are regulated by the MDF/DOT2 pathway. These transcriptome data are in agreement with RT-qPCR analysis of *PLT*, *PIN*, *SCR* and *SHR* genes in *mdf-1* (Table S1; Fig. S4G; Fig. S5; Casson et al. 2009), and further validation of the RNA-seq data by RT-qPCR was revealed as the corresponding reduction of expression of cell cycle-related genes and upregulation of stress-responsive genes (Fig. S5). The RNA-seq analysis, in the context of the phenotypic analysis, therefore suggests a role for MDF/DOT2 in promoting auxin-mediated and other signalling pathways to promote meristem function and stemness, while suppressing pathways associated with stress response and differentiation pathways.

Alternative splicing analysis was initially performed by comparing the RNA-seq data to the Arabidopsis reference transcriptome AtRTD2 (Zhang et al., 2017). This identified 4706 genes that were spliced differentially compared with wildtype with a false detection rate (FDR) of 0.01 and a 10% inclusion difference minimum (Supplementary MATS files). The most frequently detected mis-splicing events in *mdf-1* were retained introns, both increased (1028 events) or decreased (1597 events) frequencies compared with wildtype, followed by alternative 5’ splice site use (477 increased, 333 decreased events) and alternative 3’ splice site use (412 increased, 451 decreased events), skipped exons (279 increased, 110 decreased events), and least frequent were other, multiple exon events (11 increased, 8 decreased events) (Fig. S4H). Gene enrichment analysis of the 2015 alternative splicing events identified by RMATs analysis showed that the most common pathways affected were ‘mRNA metabolic process’, ‘vegetative to reproductive phase transition of meristem’ and ‘RNA splicing’ (p<0.05) (Fig. S4I). These results are consistent with MDF being required for correct gene expression control via splicing, and we aimed to identify other root meristem genes that may be spliced by a MDF-dependent mechanism.

### MDF controls meristem activity through RSZ33 and ACC1

Sixty-five alternatively spliced gene transcripts were identified associated with the ‘meristem’ GO category (Fig. S4I). One such gene is *RSZ33* (AT2G37340), which encodes a mRNA that is alternatively spliced (Palusa et al., 2007). It is strongly expressed in the root meristem (Birnbaum et al., 2003), with 4 RNA isoforms identified across leaf, root and flower (Simpson et al., 2008), of which 3 isoforms are found in the root meristem (Li et al. 2020; Fig. 2A,B). *RSZ33* encodes a plant-specific member of the SR protein family, and so is itself potentially a splicing factor (Fig. S3D). A second gene, *ACC1* (AT1G36160) is required for fatty acid biosynthesis (Baud et al. 2004) and the RNA has 3 isoforms following splicing in the 5’ UTR in different tissues (Fig. 2C-E). Interestingly, defective expression of each gene (namely in transgenic overexpressers of *RSZ33* and in the *acc1* allelic mutants *gurke* and *pasticcino3*) leads to root growth defects similar to those seen in *mdf*, such as a short root and other embryonic and postembryonic defects (Baud et al. 2004; Kalyna et al., 2003; Fig. 2F-H; = Fig. S3C). Both *RSZ33* and *ACC1* transcripts are mis-spliced in the *mdf* mutant, producing either premature termination codons that likely lead to nonsense-mediated degradation of the RNA (NMD; Kalyna et al. 2012) and observed reduced transcript levels for *RSZ33* (Fig. 2B) or an alternative 5’ splice donor site usage for *ACC1* (Fig. 2D, Supplementary MATS Files). These results show that their correct splicing is dependent on MDF activity.

**Fig. 2.**
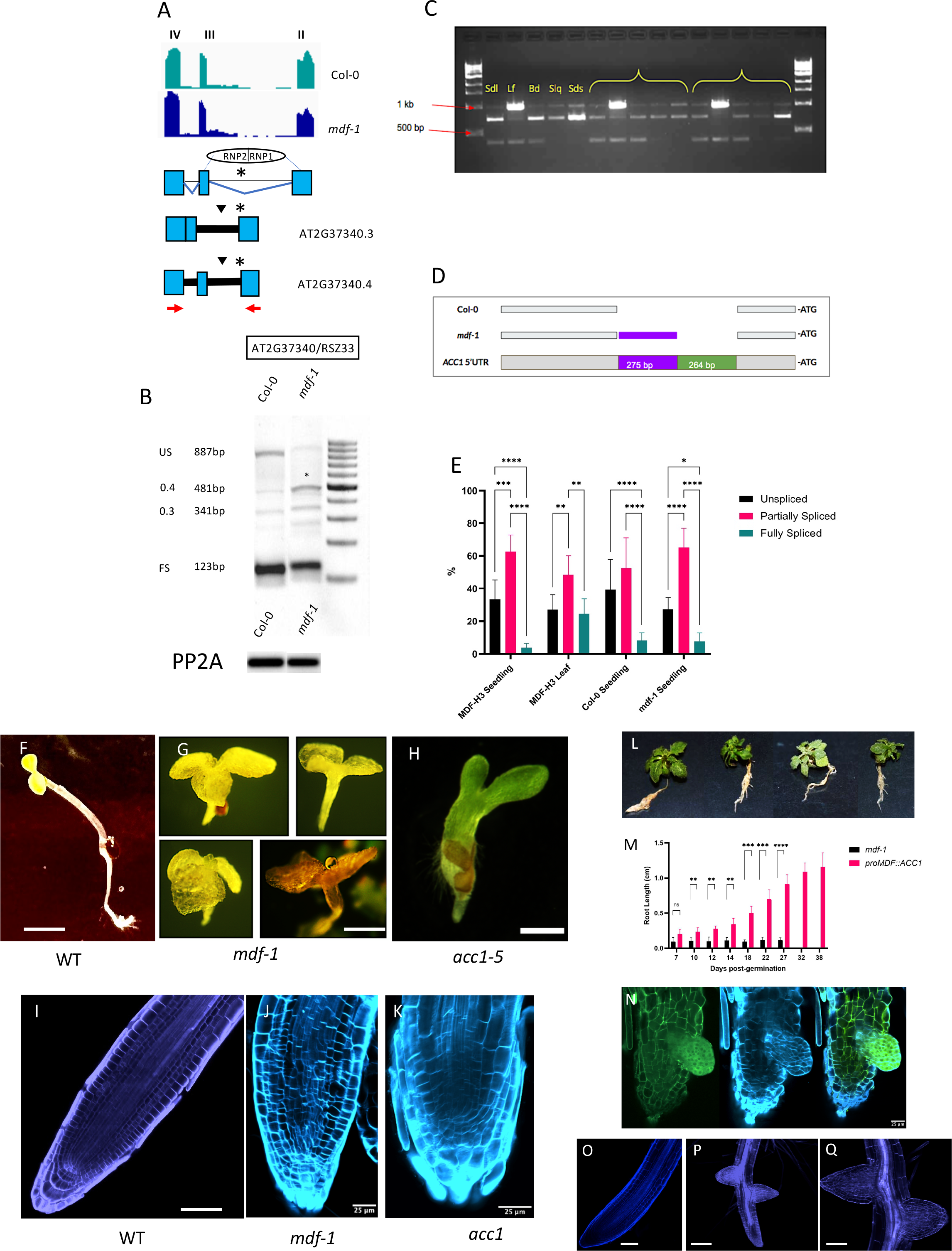
*RSZ33* and *ACC1* transcripts are mis-spliced in *mdf* and regulate. **A.** Representation of the RSZ33 A3’SS events. Inverted black triangles indicate premature termination codons. The asterix indicates the position of the alternative 3’ splice site. Red arrows indicate the primer positions used to perform the RT-PCR analysis. **B.** Root cDNA RT-PCR showing range of splice variants obtained. Shoot cDNA RT-PCR did not show any significant difference in the ratios of splice variants. In *mdf-1* root RT-PCR the RSZ33 481 bp splice variant was significantly (< 0.05) more abundant than in Col-0. The RS40 710 bp splice variant was more abundant than in wild type with a <0.001 level of significance. Control RT-PCR with PP2A primers demonstrates equivalent levels of mRNA (lower panel). **C.** Characterisation of the *ACC1* 5’UTR in 1% agarose gel by RT-PCR. Three size bands were obtained: 1kb, ca. 750 bp, and ca. 450 bp. The samples were: Seedling (Sdl), leaf (Lf), bud (Bd), silique (Slq), and seeds (Sds). Three independent biological replicates were used. **D.** Digrammatic representaiton of alternative donor site in the *ACC1* 5’UTR in wildtype (Col-0) and *mdf-1*. **E.** Abundance of *ACC1* 5’UTR splicing variants. Asterisks indicate significance levels, using ordinary two way ANOVA with Tukey’s multiple comparison test: ****p<0.0001, ***p<0.001, **p<0.01, 95% CI. **F.** Wildtype seedling at 5 dpg. Scale bar = 0.5 cm. **G.** *mdf-1* seedlings at 27 d.p.g. Scale bar = 0.5 cm. **H.** *acc1-5* seedling at 5 dpg. Scale bar = 0.25 cm. **I.** Confocal image of wild type Arabidopsis root tip at 7 dpg, stained with propidium iodide. Scale bar = 50 um. **J.** Confocal image of *mdf-1* root tip at 7 dpg, stained with calcofluor white. Scale bar = 25 um. **K.** Confocal image of *acc1-5* root tip at 7 dpg, stained with calcofluor white. Scale bar = 25 um. **L.** Transgenic *mdf-1* mutants expressing *proMDF∷ACC1:EGFP* at 38 dpg, showing phenotypic rescue. **M.** Root length comparison between *mdf-1* and *proMDF∷ACC1:EGFP* complemented seedlings. Two way ANOVA and Šídák’s post-hoc multiple comparisons test. **p<0.01, ***p<0.001, and ****p<0.0001 **N.** Confocal image of *proMDF∷ACC1:EGFP* seedling root tip at 5 dpg. Left: GFP filter. Centre: calcofluor white filter. Right: merged imaged. In green: GFP; in blue: calcofluor white. Scale bar = 25 um. **O.** Confocal image of wild type root tip at 7 dpg, stained with calcofluor white. Scale bar = 25 um. **P, Q**. Transgenic *mdf-1* seedlings expressing *proMDF∷RSZ33* at 15 dpg, showing partial rescue of primary and lateral root development.

To investigate the functional relationship between MDF, RSZ33 and ACC1, we transformed the *mdf-1* mutant with non-spliceable (i.e. correctly and fully spliced) versions of either *RSZ33* or *ACC1* under the control of the *MDF* promoter, to determine whether either gene could rescue the *mdf* mutant root phenotype. The results show that each introduced gene can partially rescue both primary and lateral root development in the *mdf-1* mutant (Fig. 2I-Q), indicating that both genes are downstream of MDF splicing control and are required for MDF function in root development.

### Splicing factor SR34 is a target of MDF and mediates specific abiotic stress responses

SR34 (AT1G02840) is a general splicing factor with at least seven different splice isoforms, resulting in protein variants differing in their RS domain (Lopato et al., 1996). SR34 is highly expressed in the root meristem and shoots during early development and vegetative growth (Lopato *et al*., 1996; Stankovic et al., 2016), and plays an active role in many post-splicing processes, including the export, stability, and translation of mRNA. In the *mdf-1* mutant, the *SR34* transcript is mis-spliced compared with wildtype, with alternative 3’ splice site selection and a retained intron event (Supplementary RMATS files). A SALK T-DNA insertion mutant *sr34* was identified and confirmed as homozygous for insertion in, for phenotypic comparison with wildtype and *mdf* under standard and abiotic stress conditions. Under control conditions, WT and *sr34* seedlings exhibited no discernable difference in primary root length (Fig. 3A). A similar result was seen with seedlings grown in the presence of 150 mM NaCl, which induces salinity stress and a shorter primary root compared to controls (Fig. 3B). However, root lengths were significantly differentially reduced when seedlings were grown on media supplemented with 300 mM mannitol and 1% sucrose, which confer osmotic stress, 0.5 μM ABA, which is a hormone signal activated upon osmotic stress (Rowe et al., 2016), showing that *sr34* mutants are more sensitive than WT to these specific conditions (Fig. 3C-E). This indicates that MDF plays a role is response to osmotic stress conditions, but not salinity stress, in part via SR34 splicing control.

**Fig. 3.**
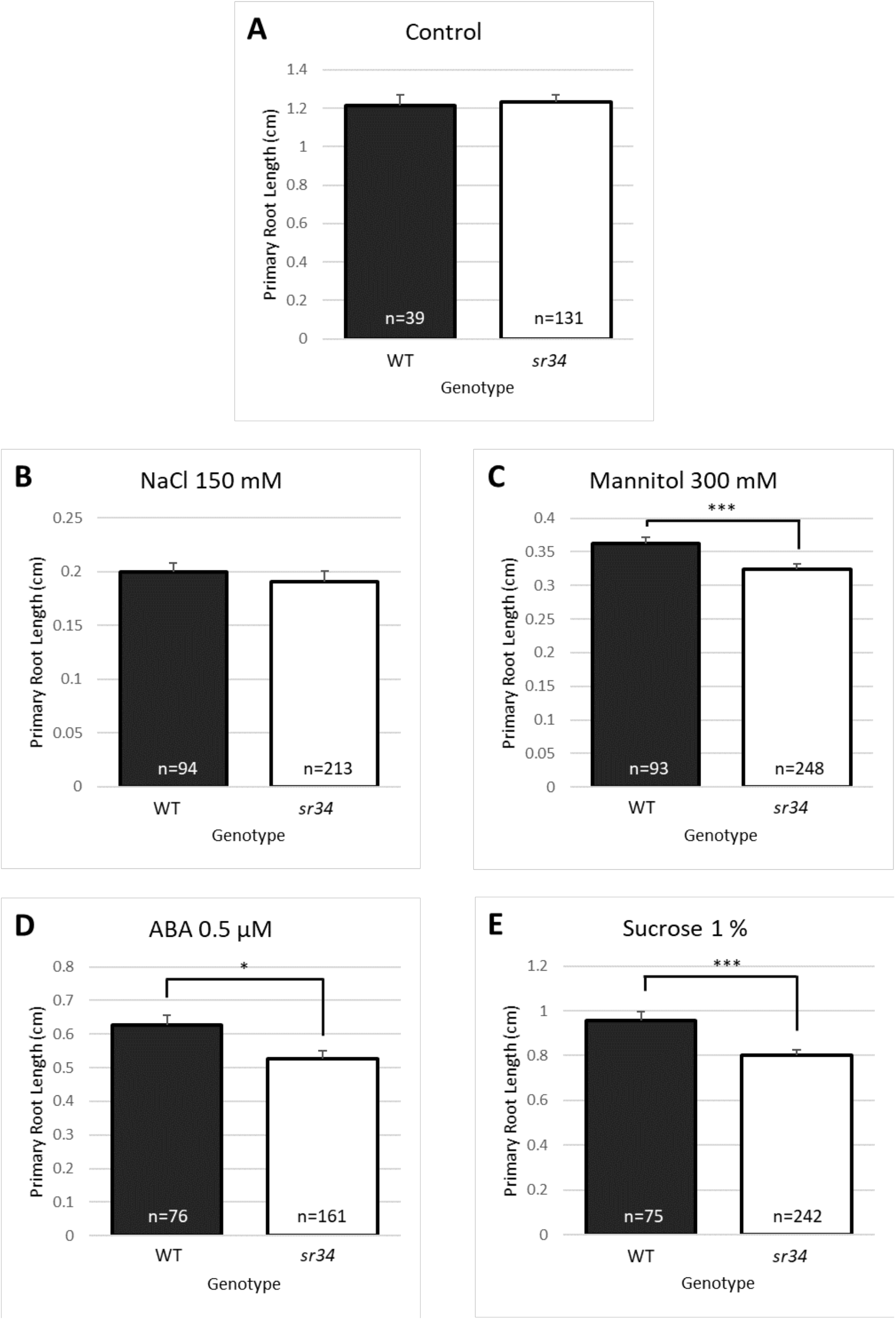
SR34 mediates specific abiotic stress responses. Primary root lengths were measured for wildtype (WT) and *sr34* mutant seedlings 7 days post germination following growth on half-strength MS medium (Control, A) or half-strength MS medium supplemented with 150 mM NaCl (B), 300 mM mannitol (C), 0.5 μM ABA (D) or 1% sucrose (E). Root length values are means of the sample sizes indicated, with the sample sizes reflecting three pooled biological replicates. Error bars represent +1 standard error. Asterisks indicate significance levels, using ordinary two way ANOVA with Tukey’s multiple comparison test: ****p<0.0001, ***p<0.001, **p<0.01, 95% CI.

### MDF RNA is spliced and isoforms are regulated by abiotic stress

Given the evidence that MDF promotes stem cell maintenance and represses cell differentiation, we hypothesized that the level of *MDF* expression may provide a mechanism to maintain meristem function under abiotic stress conditions. To test this, we determined whether *MDF* expression or splicing is modulated by environmental stresses that affect root meristem activity and growth.

Osmotic stress reduces root growth in Arabidopsis via an interacting signalling network invoving ABA, ethylene and *PIN* gene regulation that leads to low accumulated auxin in the root meristem (Rowe et al. 2016). To determine whether MDF may be part of this mechanism, we used RT-qPCR to measure *MDF* expression under osmotic stress (1.1-1.4 MPa using PEG 8000; Rowe et al., 2016) or salt stress. Results show that *MDF* transcription is induced by osmotic and salt stress within 30 min of treatment, significant at p<0.01 (Fig. 4A). Interestingly, the *MDF* transcript is itself alternatively spliced, and the levels of a 3’UTR retained intron isoform rise significantly between 6 and 12 h after osmotic shock in root tissue (Fig. 4B,C). This supports a role for MDF in maintaining meristem pathways and suppressing cell death/differentiation under stress conditions. Similarly, under cold stress, we found (using transcriptional data described in Calixto et al. 2018) there is also isoform switching, with the retained intron transcript accumulating from a low level within 30 min of treatment, while the fully spliced isoform declines (Fig 4D).

**Fig. 4.**
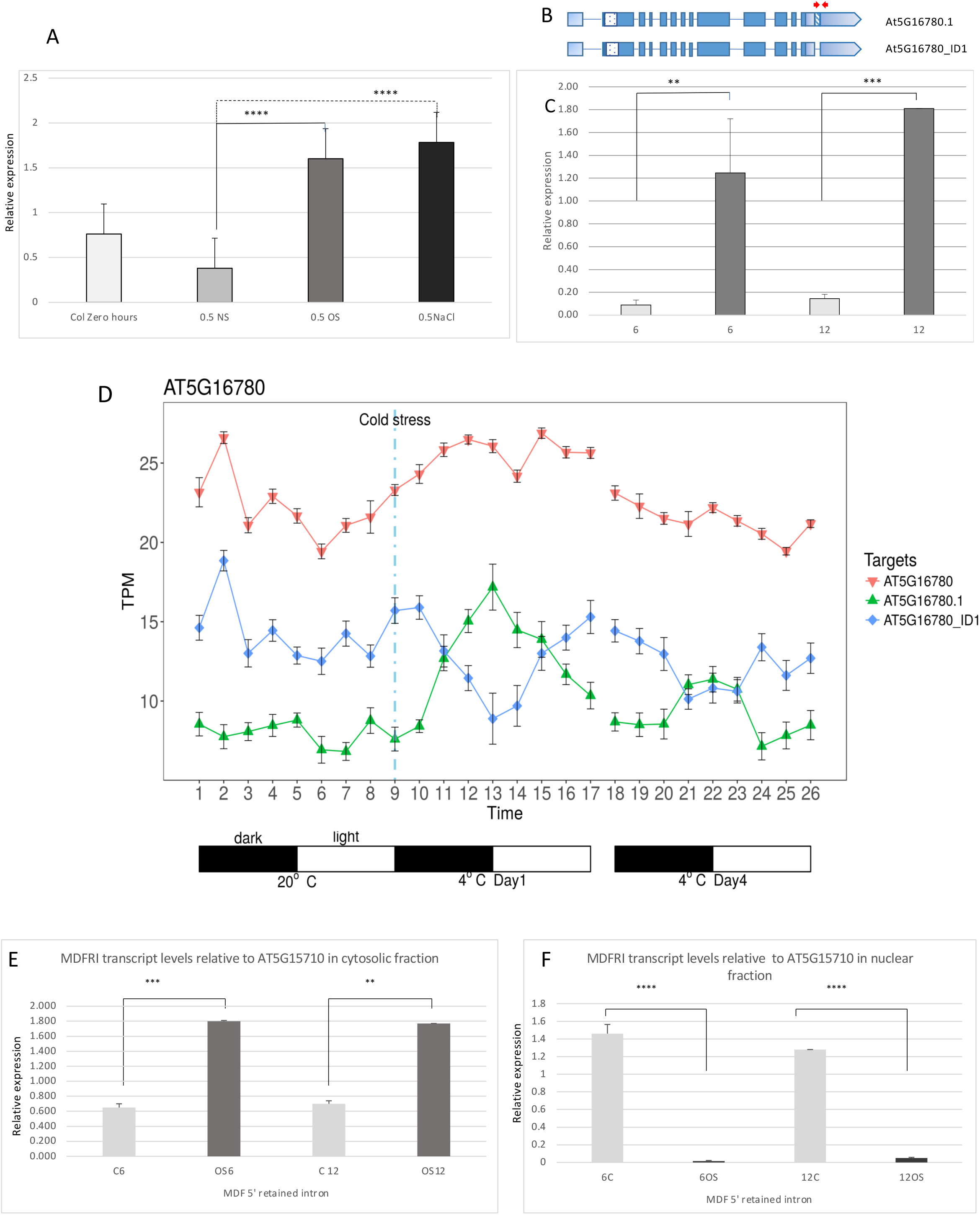
*MDF* expression in response to abiotic stress. **A.** *MDF* expression (measured by qRT-PCR, relative to *UBQ10* expression) in wild type seedling roots (Col-0) at 7 dpg, at time zero (Col 0 time zero), or transferred for 30 min to standard medium (no stress), 30 min osmotic stress (1.1-1.4 MPa using PEG 8000) or salt stress (150 mM NaCl). **B.** Gene model of *MDF*. Untranslated regions are shaded pale blue, the C-terminal RS domain is the spotted box, and the 3’ retained intron is shown by the blue diagonal lines. Red arrows represent the primers used in qRT-PCR. **C.** qRT-PCR results showing the levels of the 3’UTR retained intron in root tissue before and after osmotic stress. Levels significantly rise six and twelve hours after osmotic shock (1.1-1.4 MPa PEG 80000, dark grey bars). With no stress, the levels remain low and unchanged with time (pale grey bars). **D.** Switching of *MDF* splicing isoforms (total transcript, AT5G17780; retained intron isoform, AT5G17780.1; spliced isoform, AT5G17780_ID1) under cold stress. At the onset of cold stress (vertical broken blue line) the retained intron transcript (green line) accumulates from a low level within 30 min of treatment, while the fully spliced isoform (blue line) declines. Light-dark cycles are indicated. **E,F**. qRT-PCR analysis to determine cytosolic (d) and nuclear (e) distribution of MDF isoforms with retained intron under control (non-stress) conditions for 6 or 12 hours (C6, C12) or under osmotic stress for 6 or 12 hours (OS6, OS12). Primers MDFUTRintron5’F and MDFRIR were used to detect the un-spliced AT5G16780.1 isoform. Asterisks indicate significance levels, using ordinary two way ANOVA with Tukey’s multiple comparison test: ****p<0.0001, ***p<0.001, **p<0.01, 95% CI.

Retained introns can influence the intracellular partitioning of transcripts between nucleus and cytoplasm, as a mechanism to avoid NMD (Rausin et al. 2010; Naro et al. 2017), and intron retention is a potential mechanism to ensure transcripts are retained in the nucleus and may be released following splicing in response to environmental stress (Jia et al., 2020). To determine whether the retained intron (RI) isoform of the *MDF* transcript is differentially partitioned, we purified nuclei and cytoplasm from Arabidopsis roots, isolated RNA and quantified the RI isoform in each fraction by RT-qPCR. The results show that at both 6 h and 12 h after osmotic stress, the RI isoform was relatively abundant in the cytosol (and presumably available for translation to preserve meristem function), while in the nucleus its abundance was reduced under osmotic stress, compared to non-stress conditions (Fig. 4E,F). This suggests that there is differential partitioning of the RI isoform of *MDF* between nucleus and cytosol in response to osmotic stress, though further work is required to determine what, if any, biological role the retained intron might play.

Our results show that MDF regulates the growth of roots by controlling the splicing of regulatory mRNAs and the levels of other regulatory gene transcripts that comprise a network to determine the balance between stemness and meristem activity on the one hand, and cell division arrest and cell differentiation on the other (Fig. S6). This balance is determined by the level of MDF, which itself is regulated by stress responses that modulate RNA isoform ratios to regulate growth responses through the network.

## MATERIALS AND METHODS

### Plant material and growth conditions

T-DNA insertion mutant lines *mdf-1*, *mdf-2*, *acc1* and *sr34* were obtained from Nottingham Arabidopsis Stock Centre (NASC). Seeds were sterilised with 70% ethanol and 10 % bleach and then washed five times with sterile distilled water, followed by stratification at 4°C for 4-7 days to synchronise germination and grown on half-strength MS10 medium agar plates as described (Casson et al., 2009).

### RNA extraction and sequencing

RNA was extracted from three independent biological replicates using 7-day-old seedlings (ca. 100 mg tissue) grown on half-strength MS10 medium using the Sigma-Aldrich Plant Total RNA Kit (catalogue number STRN50), with the On-Column DNase I Digestion Set (catalog Number: DNASE10-1SET) to eliminate any residual DNA molecules. Plant tissue was ground in liquid nitrogen before incubation in a lysis solution containing 2-mercaptoethanol at 65°C for 3 minute. The solid debris was removed by centrifuging at 14 000 x *g* and column filtration before RNA was captured onto a binding column using the supplied binding solution, which helps preventing polysaccharide and genomic DNA from clogging the column. Most DNA was removed by wash solutions, and any trace of residual DNA was removed by DNase on the column. Then purified RNA was eluted using RNAase-free water.

RNA sequencing from three biological replicate samples was carried out on an Illumina HiSeq 2500 System with the library prepared using the Illumina TruSeq Stranded Total RNA with Ribo-Zero Plant Sample Preparation kit (catalog Number: RS-122-2401). Ribosomal RNA (rRNA) was removed from isolated total RNA using biotinylated, target-specific oligos on rRNA removal beads. Purified RNA was quality checked using a TapeStation 2200 (Agilent Technology) with High Sensitivity RNA ScreenTape (catalogue number 5067-5579), and the mRNA was fragmented into 120-200 bp sequences with a median size of 150 bp. Fragmented mRNA was used as a template to synthesise first-strand cDNA using reverse transcriptase and random primers, followed by second-strand cDNA synthesis with DNA Polymerase I and RNase H. Newly synthesised cDNA had a single adenine base added with ligation of adaptors, before being purified and amplified by PCR to make the final library. Library quality control was performed again using a TapeStation with D1000 ScreenTape (catalogue number 5067-5582).

### Pre-processing of RNA-seq data, differential expression and differential usage analysis

RNAseq data were processed and aligned against the TAIR10 (EnsemblePlants) genome using TopHat and indexed with Samtools. DeSeq determined differential expression. Alternative splicing analysis was determined using RMats (*p* value of 0.05, a minimum of 10% inclusion difference). Alternative splicing events were visualised using Sashimi plots generated by the Integrative Genomics Viewer (IGV; Robinson et al., 2011).

### Direct mRNA isolation and cDNA preparation for RT-qPCR or RT-PCR

Seedlings were grown 7 days post-germination as described above. Roots and cotyledons were separated using a razor blade, and the material was frozen immediately in liquid nitrogen. Pools of seedlings were used to generate three separate biological samples. Each pool contained approximately 20 mg of root or cotyledon tissue. Total mRNA was extracted using Dynabeads®mRNA DIRECT™kit with Oligo(dT)_25_ labelled magnetic beads. Frozen tissue was ground with a sterile plastic micropestle and resuspended in 300 μl lysis buffer. The solution was then forced through a 21-gauge needle in a 1ml syringe 3-5 times to shear any DNA and mixed with 50 μl of Dynabeads Oligo(dT)_25_. The kit procedure was followed, with two final washes conducted. To ensure the complete removal of any genomic DNA in the subcellular fractionation experiments, this stage was followed by ezDnase™ treatment in a 10 μl volume (1μl ezDNASe™, 1 μl ezDNASe™ 10X buffer and 8 μl sterile H_2_O), 37°C for 2 min followed by 1 μl DTT and 5 min at 55°C in a heat block.

cDNA was prepared using a SuperScript®IV First-Strand synthesis system directly on the bead solution. For RT-PCR and RT-qPCR beads were washed in 20 μl 1 x SSIV buffer before resuspension in 12 μl sterile H_2_O with 1 μl dNTP 10 mM each mix and incubated for 5 min at 50 °C in a Proflex PCR machine (Applied Biosystems). Then the following were added 4 μl 10 x SSIV buffer, 1 μl ribonuclease inhibitor and 1 μl Superscript®IV reverse transcriptase were added. The mixture was mixed by pipetting and incubated for 10 min at 50 °C, followed by 10 min at 80 °C, and then held at 4 °C. The 20 μl cDNA mix was stored at −20 and not eluted from the beads.

Samples were checked for the presence of genomic DNA by PCR with Actin 2 primers ACT2 forward and reverse. A PCR reaction after 28 cycles with a Tm of 60 °C generated a 340 bp product if genomic DNA was a contaminant, 240 bp otherwise. All PCR and sequencing primers are listed in Table S3.

### RT-PCR

0.5 – 1 μl of cDNA/bead mix were used per PCR reaction. RT-PCR was performed with *RSZ33* or *ACC1* root-derived cDNA using Phusion™ (Thermofisher) high-fidelity polymerase. Relative levels of *RSZ33* and *ACC1* splice variants were determined using FIJI gel analysis software (Schindelin et al. 2012). Relative levels of cDNA per sample were determined using PCR-amplified *PP2A* transcript levels.

### Subcellular fractionation

The Intact line UBQ10∷NTF (Marques-Bueno et al., 2016), expressing NTF in the QC of the root meristem, was supplied by Nottingham *Arabidopsis* Stock Centre (NASC) and used to generate a bulk stock. Seeds were sterilised, stratified and sown on 0.5 MS and harvested at ten days. Roots were cut from the cotyledons using a razor blade, and biological replica pools generated from 20 - 25mg of root tissue. Sub-cellular fractionation was performed based on the method of Hartmann et al. (2018). The final nuclear fraction was resuspended in 200 μl HONDA buffer and, along with 200 μl of cytosolic fraction, were mixed with an equal volume of lysis buffer and 50 μl of Dynabeads® labelled with Oligo(dT)_25_. Prior to making the cDNA, the mRNA-bound bead solution was treated with ezDnase™ as described above to remove any trace of genomic DNA.

### PCR with osmotic shocked root subcellular fractions

Quality control PCR was carried out on 1μl of cDNA bound to Dynabeads®. To identify genomic DNA contamination, Act2 primers were used; a 240 bp band present indicated no contamination. MDF5’UTR and MDFRIR primers amplified the retained intron (120 bp product). Products were run on a 1% agarose gel with a 100 bp size ladder. Phusion™ (Thermofisher) high fidelity polymerase was used.

### qRT-PCR following osmotic shock

10 day-old seedlings were placed on 0.5 MS agar plates treated to achieve 1.1-1.4 MPa PEG 80000 using the method of Rowe et. al (2016). Seedlings were transferred via sterile nylon mesh to reduce any physical shock to the roots. Strips of seedling-bearing mesh were moved from untreated 0.5 MS plates onto osmotic shock and untreated control plates and returned to the growth chamber for 6 and 12 hours. A zero-hour set of three biological replicates were harvested to ensure no effect was caused by moving the seedlings. At the determined time, seedlings were harvested, and roots and shoots were separated in the case of the Intact UBQ10:NTF seedlings. Tissue was transferred to sterile Eppendorf tubes and frozen in liquid nitrogen. mRNA and cDNA were prepared using the Dynabeads™ protocol.

qRT-PCR was conducted using three biological replicates and three technical replicates for each sample. PCR Biosystems PCRBIO™ SYBrgreen was used as per the manufacturer’s instructions. Relative expression levels were determined using the ΔΔCT method relative to expression of a paired reference gene amplification. The reference gene *AT5G15710* was used due to its stable expression pattern under osmotic stress and at various developmental stages (Czechowski *et al*., 2005). Primers are listed in Table S3.

### MDF Tertiary Structure Comparative Modelling

A comparative model of MDF was generated by MODELLER (https://salilab.org/modeller/) and superimposed on SART1 in the human B complex, and the secondary structure prediction for the aligned sequences of MDF1, human SART1 (5o9z_P) and yeast Snu66 (5NRL_E). MDF template structures were searched using GeneSilico (http://genesilico.pl/meta) and the top-ranked available templates, sections of human SART1 or yeast Snu66 were used to model MDF. The modelled C-terminal helix of MDF was removed for Figure 2.

### Bimolecular fluorescence complementation (BiFC)

Fully spliced versions of MDF, BRR2A, PRP6 and PRP8 were subcloned into Gateway ™ vectors pDNOR207 or pDONRZeo and subsequently into both BiFC plasmid vectors pYFN43 and pYFC43 (gift of Dr. Patrick Duckney, Durham University) using the Gateway ™ Clonase system. Positive clones were identified by colony PCR and then DNA sequenced prior to use in BiFC interactions. Positive controls were pYFC43BIN2 and pYFN43BZR1 (gift of Dr. Miguel de Lucas, Durham University).

BiFC constructs (500 μg) were transfected into GV3101 *Agrobacterium tumefaciens* cells and selected with 30 μg/ml gentamycin, 50 μg/ml kanamycin and 50 μg/ml rifampicin. Single colonies were selected and grown then diluted to an OD600 0.2 in LB with 200 μM acetosyringone. GV3101 colonies transformed previously with a p19 construct to suppress gene silencing were also grown. After growth to OD600 0.4-0.6 cells were resuspended into MMA buffer (MS 5 g/l, MES 1.95 g/l, sucrose 20 g/l, pH adjusted to 5.6 using NaOH) plus 200 μM acetosyringone, and shaken for 1 hour at room temperature in the dark. The OD600 determined volumes used combined cultures pYFN*x* + pYFC*x*+ p19 in a ratio of 1:1:1 in 1ml volume.

### Agroinfection and transient expression in tobacco leaf

*Nicotiana benthamiana* plants were grown for 3 weeks at 21°C with 16-hlight/8-h dark cycles. Mixed cultures were slowly infiltrated using a 1 ml syringe into the lower leaf epidermal layer until the whole leaf was saturated. Plants were covered and left in the subdued lighting overnight before being returned to the growth cabinet for 3 days. 0.5 cm sections were taken for microscopic analysis. Imaging of fluorescence in leaf epidermis was as described previously (Gu et al., 2021).

### Gateway™ Cloning

#### BP reaction

From the 5’ upstream region 1075bp of the *MDF* promoter and 873bp of the fully spliced *RSZ33* and *ACC1* coding sequences were synthesised into an artificial fusion within the pEX-A258 vector (Eurofins) with the Gateway ™ attB1 and attB2 sequences at the 5 ’and 3’ ends. Two sets of primers were designed to amplify the *MDF* promoter and the *pDONR207∷ ACC1* backbone from the *pDONR207∷ACC1* entry clone (Table S3). The resulting clones were sequenced to ensure no mutations or errors were present. The attB linkers then provided the means to transfer the construct into the Gateway ™ entry vector pDONR207 via BP reaction. In a 5 μl reaction, 100 ng of donor vector (pEX-A258-MDFPromRSZ33), 100 ng pDONR207 plasmid, 2 μl of TE buffer pH8 and 1 μl Gateway ™ BP Clonase II enzyme (Thermofisher) were combined in a 20 μl PCR tube. After mixing by pipetting the reaction was placed at 25 °C in a heat block overnight. To stop further reaction 1 μl of proteinase K was added and the reaction tube was heated to 37 °C for 10 minutes. 2.5 μl of the reaction mix was transformed into Sub-cloning efficiency ™ DH5α competent cells (Thermofisher) and plated onto Lagar with 50 μg/ml Ampicillin selection.

#### Colony PCR

After growth overnight at 37°C, colony PCR was performed to identify positive colonies. Gene-specific PCR primers (Table S3) were used in a reaction with Dreamtaq (Thermofisher). Colonies were picked into 20 μl sterile water with a micropipette tip and resuspended. The suspension was also streaked to single colonies on a fresh L-Gentamycin 10 μg/m plate. One μl of colony suspension was used per PCR reaction with 1 X buffer, 200 μM each dNTP, 0.5 μM of each primer, 1.25 U Dreamtaq polymerase in a 50 μl total reaction volume. Reactions were performed at 95 °C for 5 min, then 25 cycles of 95°C 30 s, 56 °C 30 s, 72 °C 30 s, followed by 1 cycle 5 min at 72 °C. 1 ng of pEX-A258-MDFPromRSZ33/ACC1 were used as controls. PCR products were analysed on a 1% agarose gel with 100 bp markers.

#### DNA Sequencing

PCR-positive clones were isolated and each cultured to prepare plasmid DNA for sequencing. Construct fidelity was confirmed by the Sequencing Service, Dept. of Biosciences, Durham University. DNA sequences were analysed using SnapGene software.

#### LR reaction and GV3101 transformation

100 ng of the vector pMDC107_B with Basta resistance gene (gift of Dr. J. Kroon, Durham University) was used as the destination vector for the Gateway ™ LR reaction containing 100 ng of pDONR207MDFprom∷RSZ33, 3μl TE pH 8 buffer and 1 μl Gateway ™ LR Clonase II enzyme (Thermofisher). The reaction was mixed and incubated at 22 °C for 3 h, followed by the addition of 1 μl proteinase K and a further incubation at 37 °C for 10 min. The whole reaction was used to transform MAX efficiency™ DH5α competent cells (Thermofisher) and selected on L agar + kanamycin 50 μg/ml. To produce the *proMDF∷ACC1:EGFP* expression clone, purified *proMDF∷ACC1* entry clone was used for the LR reaction, cloned into the pJK1243 destination vector, and introduced into chemically competent DH5α cells. Positive colonies were confirmed by PCR. Sequencing of the expression clone was done only to ensure that the protein fusion between *ACC1* and *GFP* DNA sequences was in the correct reading frame.

Plasmid DNA was prepared from a positive clone, and 1 μg was used to transform the *Agrobacterium tumefaciens* strain GV3101 (lab stocks), a negative H_2_O control was included. Positive transformants were selected on gentamycin 30 μg/ml, kanamycin 50 μg/ml and rifampicin 50 μg/ml. Arabidopsis was transformed with the respective *proMDF∷RSZ33* and *proMDF∷ACC1:EGFP* constructs by the floral dip method (Clough and Bent, 1998).

#### *mdf-1* complementation by transformation

To determine whether non-spliceable versions of *RSZ33* or *ACC1* could rescue the *mdf-1* mutant, heterozygous *mdf-1* mutants were used to generate flowering plants for transformation, as homozygote mutants do not flower. Six plants per dipping were transformed by floral dip (Clough & Bent, 1998) using Agrobacterium containing either *proMDF∷RSZ33* or *proMDF∷ACC1:GFP* (fully spliced variants). Seeds from several individual transformation events were collected after 3-5 weeks when the siliques were brown and dry, for selfing to generate mutants homozygous for *mdf-1* and containing either the *proMDF∷RSZ33* or *proMDF∷ACC1:GFP* transgenes, for phenotypic analysis.

#### Selection of transformants

Stratified T1 seeds (2 days at 4°C) from independent plants were sown at high density on compost and grown at 22 °C (c.3000 lux) in a growth chamber (Sanyo Electric) in a 16 h photoperiod. After 10 days putative *proMDF∷RSZ33* x *mdf-1* seedlings were sprayed 3 times with 250 mg/l commercial Basta (Kurtail Evo) at 2 day intervals (Harrison et al., 2006). Surviving transformants were transferred to individual pots and allowed to flower. T2 seed was collected in a covering seed bag to prevent cross-contamination. Once brown and dry the siliques were harvested and stored.

Transformation was confirmed by genotyping. 10 day-old cotyledons were used in two PCR reactions using the MyTaq™ Plant – PCR kit (Meridian Biosciences®). 2 x 2 mm slices of cotyledon were added directly to a 50 μl reaction. Two separate PCR reactions were performed to identify the construct with relevant primers to determine whether the *MDF* promoter is fused to the *RSZ33* or *ACC1* coding sequences.

To identify any developmental effect of the *MDF* promoter coupled to non-spliceable versions of *RSZ33* or *ACC1* on *mdf-1* phenotype, 10 day-old seedlings were selected. *proMDF∷RSZ33* x *mdf-1* seeds were sown onto medium containing 50 mM phosphinothricin solution (Basta active ingredient; Sigma-Aldrich) in 0.1% agar (Harrison et al., 2006). *proMDF∷ACC1:GFP* x *mdf-1* homozygous mutant plants were selected by phenotype, GFP fluorescence and PCR. Seedlings with the typical *mdf-1* phenotype were transferred to a new plate to be observed under the fluorescent stereo microscope (Leica M165 FC Fluorescent stereo microscope equipped with a Leica DFC 420C camera) before confocal imaging.

#### Root tissue preparation and microscopy

Separated roots were individually placed in a sterile plastic well plate and then fixed in 4% paraformaldehyde for 30 min under vacuum. Roots were then gently washed twice in 1 x PBS. Samples were stored in ClearSee solution (Kurihara et al., 2015) for a minimum of 4 days prior to observation by microscopy. Control wild-type root tissue was prepared for comparison. For seedlings stained with Calcofluor White, 0.1% Calcofluor White in ClearSee solution was prepared and the seedlings were stained for 30 min with gentle shaking. Seedlings were washed in ClearSee for 30 min prior to imaging. Imaging used a Zeiss 800 Laser Scanning Confocal Microscope x20 objective with excitation at 405 nm.

## Author contributions

KL, JT and HT planned the research; HT, WS, RM, MK, CJ, DD, XY, NZ and SM-J carried out experimental work; KL, SG, CC, JT and XZ had supervisory roles; KL drafted the manuscript and all authors had the opportunity to edit the manuscript.

## Data availability

Raw data for RNA seq and splicing analysis (MATS) and qRT-PCR have been deposited in the Dryad Digital Repository (https://doi.org/10.5061/dryad.b2rbnzskc).

## Supplementary Data legends

**Fig. S1. The *MDF* gene and protein**

**A.** Organization of the *MDF* gene (AT5G16780) showing sites of mutations (*mdf-1*, *mdf-2* - black triangles) and expression of the proMDF∷GUS gene fusion in the root tips of transgenic Arabidopsis at 3 and 7 dpg.

**B.** Location of PCR primer sites (upper panel) and amplification of transcript in wild-type (Col-0), transgenic MDF overexpressing (35S∷MDF), *mdf-1* and *mdf-2* mutant and *mdf-1* mutant complemented with a proMDF∷MDF gene fusion. Some fragments of *mdf-2* in particular are present, consistent with some transcript detected in RNA-seq data (Table S1), but it is expected there is incomplete protein production in *mdf-2*.

**C.** SSDB protein motif search for MDF, showing strong simillarity to SART1 family proteins.

**D.** Amino acid comparison between MDF and human SART-1 protein.

**E.** Amino acid sequence of MDF predicted RNA/DNA binding domain.

**F.** Predicted helical (red) and strand (blue) regions in MDF (model), hSART-1 (509Z_P) and SnU66 (5NRL_E) proteins.

**Fig. S2 Predicted MDF structure and protein interactions in the plant spliceosome**.

A. Comparative model of the putative 3-dimensional structure of the MDF/DOT2 protein (red) suggests a strong structural similarity with hSART-1 (yellow). Predicted relationships with PRPF6 and PRPF8 (cyan) is shown.

B-I. Bimolecular fluorescence complementation (BiFC) following transient gene expression in *Nicotiana bethamiana* leaves shows interaction between BIN2 and BZR1 (positive control, B), BIN2 and MDF (C), BZR1 and MDF (D) BRR2a and MDF (F, G) but not between MDF and PRP8 (E) or between BRR2a and BZR1 (H) or BRR2a and BIN2 (I).

**Fig. S3 *mdf* mutants and *RSZ33* overexpressers have similar phenotypes**

Phenotypes of (A) *mdf-1* and *mdf-2* seedlings at 7 dpg, (B) wild-type (Col-0), *mdf-1* and *mdf-2* mutant and *mdf-1* mutant complemented plants at 4 weeks post-germination; and (C) *RSZ33* transgenic overexpressers at 8 dpg. Scale bar = 0.5 cm.

D. RSZ33 protein sequence showing zinc finger and RS domains, and site of retained intron that creates a premature termination codon (PTC).

**Fig. S4 Transcriptional analysis of *mdf* mutants**

**A**. Venn diagram showing the number of differentially expressed genes (DEGs) in *mdf-1* and *mdf-2* homozygous transgenic 7 d.a.g. seedlings following the RNA sequencing experiment, with adjusted p value < 0.05, and log2 fold change (log2fc) >1. Each oval contains all up- or down-regulated genes in one of the genotypes, and the overlapping parts represent numbers of genes meeting the conditions of more than one encircling oval. Percentage under each number is calculated by dividing each number by the total number of DEGs in the diagram.

**B-E**. Treemaps of enriched GO terms in significantly up- or down-regulated genes from *mdf-1* and *mdf-2* mutants.

**F.** Differential expression analysis (DeSeq2) of stress response genes (as revealed as GO terms) in *mdf-1* and *mdf-2* compared to Col-0.

**G.** Differential expression analysis (DeSeq2) of genes required for root meristem development and auxin transport in *mdf-1* compared to Col-0. Except for *PLETHORA5* (*PLT5*), all are down-regulated in *mdf-1*.

**H.** Comparison of the alternative splicing events between Col-0 and *mdf-1*, identified by RMATs analysis of the RNAseq data. The events compared were SE (skipped exon), RI (retained intron), MXE (multiple exon events), A5SS (alternative 5’ splice sites) and A3SS (alternative 3’ splice sites).

**I.** Gene enrichment analysis dotplot of the 2015 alternative splicing events identified by RMATs analysis with a p-value < 0.01 and minimum ± 10 % inclusion difference. The most represented GO term, the largest dot, contained mRNA metabolic process candidate genes followed by vegetative to reproductive phase transition of meristem.

**Fig. S5 Expression of selected meristem and stress-related genes in *mdf* mutants and *MDF* overexpressers**

qRT-PCR analysis of selected genes involved in meristem function, auxin transport and response, U SnRNP RNA processing, cell cycle and stress response in wild-type (Col-0), transgenic MDF over expressing (35S∷MDF), *mdf-1* and *mdf-2* mutant and *mdf-1* mutant complemented with a proMDF∷MDF gene fusion at 7 dpg.

**Fig. S6 Role of MDF in control of stemness and cell differentiation**

Model describing the relationship between MDF and the antagonistic processes of stem cell maintenance and differentiation, stress and cell death. Functional levels of MDF increase under stress to suppress cell death and differentiation, and to maintain stem cell identity and cell division activity in the meristem.

**Table S1**

RNA-seq data showing up- and down-regulated expression levels of selected meristem genes of interest for *mdf-1* and *mdf-2*.

**Table S2**

AtRTD2 DeqSeq data showing fold-change transcript levels in *mdf-1* compared to wildtype at fdr of 0.05.

**Supplementary MATS Files: Multivariate analysis of transcript splicing (MATS)**

Files listing counts of differential alternative splicing events recorded between wildtype and in the *mdf-1* mutant. The events compared were skipped exon (SE), retained intron (RI), multiple exon events (MXE), alternative 5’ splice sites (A5SS) and alternative 3’ splice sites (A3SS). Uploaded to dryad database.

**Supplementary qRT-PCR raw data**

Files with data for gene expression as described in Fig. S5.

